# *Cryptococcus neoformans* melanization incorporates multiple catecholamines to produce polytypic melanin

**DOI:** 10.1101/2021.08.26.457838

**Authors:** Rosanna P. Baker, Christine Chrissian, Ruth E. Stark, Arturo Casadevall

**Affiliations:** Department of Molecular Microbiology and Immunology, Johns Hopkins Bloomberg School of Public Health, Johns Hopkins University, Baltimore, MD 21205, USA; Department of Chemistry and Biochemistry, The City College of New York and CUNY Institute for Macromolecular Assemblies, New York, NY 10031, USA; Ph.D. Program in Biochemistry, The Graduate Center of the City University of New York, New York, NY, USA; Ph.D. Program in Chemistry, The Graduate Center of the City University of New York, New York, NY, USA

**Keywords:** fungi, virulence factor, melanin, *Cryptococcus neoformans*, cell wall, catecholamine, dopamine, solid-state NMR, reactive oxygen species

## Abstract

Melanin is a major virulence factor in pathogenic fungi that enhances the ability of fungal cells to resist immune clearance. *Cryptococcus neoformans* is an important human pathogenic fungus that synthesizes melanin from exogenous tissue catecholamine precursors during infection, but the type of melanin made in cryptococcal meningoencephalitis is unknown. We analyzed the efficacy of various catecholamines found in brain tissue in supporting melanization using animal brain tissue and synthetic catecholamine mixtures reflecting brain tissue proportions. Solid-state NMR spectra of the melanin pigment produced from such mixtures yielded more melanin than expected if only the preferred constituent dopamine had been incorporated, suggesting uptake of additional catecholamines. Probing the biosynthesis of melanin using radiolabeled catecholamines revealed that *C. neoformans* melanization simultaneously incorporated more than one catecholamine, implying that the pigment was polytypic in nature. Nonetheless, melanin derived from individual or mixed catecholamines had comparable ability to protect *C. neoformans* against ultraviolet light and oxidants. Our results indicate that melanin produced during infection differs depending on the catecholamine composition of tissue and that melanin pigment synthesized *in vivo* is likely to accrue from the polymerization of a mixture of precursors. From a practical standpoint our results strongly suggest that using dopamine as a polymerization precursor is capable of producing melanin pigment comparable to that produced during infection. On a more fundamental level our findings uncover additional structural complexity for natural cryptococcal melanin by demonstrating that pigment produced during human infection is likely to be composed of polymerized moieties derived from chemically different precursors.

## Introduction

*Cryptococcus neoformans* is an encapsulated, free-living fungal organism that can establish opportunistic infections in both plant and animal hosts (1). *C. neoformans* is broadly prevalent in the environment and colonizes a variety of ecological niches, which notably include soil and bird guano (2). Despite ubiquitous exposure of the human population, *C. neoformans* rarely causes disease in individuals with intact immunity as the infection is either cleared or becomes asymptomatically latent (3). However, *C. neoformans* poses a serious threat to those with compromised immunity resulting from disease or medical treatment, especially AIDS patients, and is estimated to cause 180,000 deaths annually (4). Cryptococcal infection commences with the inhalation of desiccated fungal cells or spores that become deposited in the lungs, which can result in cryptococcal pneumonia for immunocompromised individuals (5). Progression of the primary infection can occur through extrapulmonary dissemination as yeast cells dispersed through the bloodstream are able to cross the blood brain barrier and take up residence in brain tissue (6, 7). The ensuing meningoencephalitis that is typical of this aggressive stage of cryptococcosis, if left untreated, is almost always fatal (8). Although concerted efforts are underway to develop an effective vaccine (9), there is currently no reliable means of preventing infection and even the most current treatment strategies that employ a combined anti-fungal regimen only decrease 10-week mortality rates to 24% (10). Thus, an improved understanding of factors that contribute to fungal virulence is vital to our efforts in combating this harmful pathogen.

Among the adaptations that safeguard *C. neoformans* from harsh conditions both in the environment and during host infection is the deposition of a layer of melanin in the fungal cell wall. Melanins are a diverse group of pigments, typically black or dark brown in color, that are formed by the oxidative polymerization of phenolic or indolic precursors (11). They are notorious for their extreme recalcitrance, structural complexity, and unique biophysical properties, such as the ability to absorb a wide spectrum of UV light. Melanin pigments are widely employed throughout the biosphere, serving such diverse functions as camouflage, thermal modulation, and protection from radiation (12); in pathogenic fungi, they are linked to increased virulence (13) and drug resistance (14). In *C. neoformans* cells, melanin is synthesized from phenolic compounds in small vesicles called melanosomes that are transported and subsequently deposited within the cell wall (15). There, the melanin particles assemble into larger granules that become tightly interwoven with the constituents comprising the cell wall, which consequently thickens over time (16). In the environment, melanization is credited with protecting *C. neoformans* from electromagnetic radiation, temperature stresses, and amoeba (17, 18). During infection, melanization helps shield *C. neoformans* cells from engulfment and killing by host macrophages (19–21) and contributes to the severity of infection by promoting neurotropism (22, 23). As melanization is one of two primary virulence factors for *C. neoformans*, the disruption of this process, perhaps by inhibiting the laccase enzyme responsible for catalyzing melanin synthesis, has been widely recognized as a potential therapeutic target (24).

A defining feature of the *C. neoformans* laccase enzyme is its inability to produce melanin pigments from endogenously synthesized compounds like tyrosine, the precursor most commonly used by other melanotic organisms (25). Instead, *C. neoformans* draws on exogenous sources of substrates, producing melanin pigments from a wide variety of phenolic compounds found in natural habitats and in animal hosts, as evidenced by the isolation of melanized cells from both natural isolates (26) and infected host tissues (27, 28). Indeed, the animal nervous system serves as a rich source of precursors in the form of catecholamines, nitrogen-containing diphenolic compounds which include the neurotransmitters dopamine, epinephrine, and norepinephrine. *C. neoformans* melanization has been studied *in vitro* by culturing cells in media supplemented with each of these compounds and more commonly, with L-DOPA, the immediate biosynthetic precursor of dopamine. Considerable insight into the properties of *C. neoformans* melanin has been gained by applying biophysical techniques like solid-state nuclear magnetic resonance (NMR) and electron paramagnetic resonance (EPR) to analyze melanin containing particles derived from such cultures (29–31). These analyses have indicated that the pigments produced by *C. neoformans* from each of these precursors are indeed melanins. They have also revealed distinct differences in molecular structure and EPR signal intensity, but the implications of these differences with respect to virulence remain unclear.

Whereas previous studies of *C. neoformans* melanin have largely employed L-DOPA as the precursor (reviewed in (32)), the low abundance of this compound in host tissues prompted us to question how faithfully L-DOPA melanin produced in culture reflects the structure and function of ‘natural’ melanin produced during the course of *C. neoformans* infection. The progression of cryptococcal disease ultimately results in brain dissemination, which provides a rich and varied source of catecholamines, thereby presenting the potential for simultaneous incorporation of multiple precursors. Although acid-resistant melanin particles can be isolated from infected human and animal tissue (28, 33), the limited quantity of melanized material that can be recovered from animal models of infection has precluded extensive biophysical analyses. To circumvent this obstacle, we sought to develop a cell culture system that would approximate the repertoire of catecholamine precursors present in the human brain. Drawing on previous neurochemical studies that quantified catecholamines isolated from postmortem human brains (34, 35), we melanized *C. neoformans* in a mixture of three catecholamines according to their relative brain proportions of 60% dopamine, 33% norepinephrine, and 7% epinephrine. Here we describe the isolation and analysis of melanin produced from this brain catecholamine mixture (BCM) and provide evidence that *C. neoformans* can incorporate multiple precursors simultaneously during melanin synthesis. BCM is found to be comparable to L-DOPA melanin in its ability to protect *C. neoformans* from sources of cellular damage like UV radiation and free oxygen radicals. Despite these functional similarities, the ability of *C. neoformans* to utilize a mixture of chemically distinct precursors during melanin synthesis implies architectural heterogeneity for this polymer depending on the local abundance of precursors and significantly complicates its structural characterization.

## Results

### C. neoformans KN99α variant achieves robust melanization in conditions mimicking the human brain environment

We sought to approximate the melanization process during infection by growing *C. neoformans* in a mixture of catecholamines representative of human brain proportions (brain catecholamine mixture, BCM). A key consideration in this effort was the selection of a laboratory strain variant that would be capable of robust melanin production from BCM at the human body temperature of 37°C. Several variants derived from the widely used H99 strain of *C. neoformans* var. *grubii* (H99O), including H99E, H99C, and H99W, display reduced virulence in mice and decreased melanization on L-DOPA agar as a result of laboratory passage (36). Conversely, variants H99S, H99F, and KN99α/a that were derived from passage of H99 through the rabbit central nervous system model are more virulent and produce more melanin when grown on L-DOPA (36). We compared melanization by H99O and KN99α on agar supplemented with BCM as well as the individual precursors DA, NE, and L-DOPA at 30°C and 37°C (Fig. 1A). The well-characterized H99-derived ΔLAC1 strain deficient in Laccase-1, the primary enzyme required for melanization, was included as a negative control. *C. neoformans* laccase production and activity has been shown previously to decrease with increasing temperature (37, 38). Accordingly, we observed a slower rate of melanin production and a diminished amount of overall pigmentation for both H99O and KN99α grown at 37°C compared to 30°C, regardless of which precursors were provided. Whereas H99O melanization was delayed compared to that of KN99α at 30°C, similar degrees of pigmentation were observed for the two strains after incubation for 48 h (Fig. 1A, left panel). However, H99O cells showed significantly less pigmentation than KN99α after 48 h at 37°C (Fig. 1A, right panel; Fig. 1B) and only achieved equivalent melanization levels after an additional two days of growth (data not shown). Melanization by KN99α on BCM-supplemented agar was comparable to the pigmentation observed with each of the individual catecholamines at 30°C (Fig. 1C, left graph) and was more efficient than NE alone at 37°C (Fig. 1C, right graph). Given the increased melanin production of KN99α compared to H99O when grown in the presence of BCM at the human body temperature of 37°C, the KN99α variant of *C. neoformans* was chosen for all subsequent analyses.

**Figure 1.**
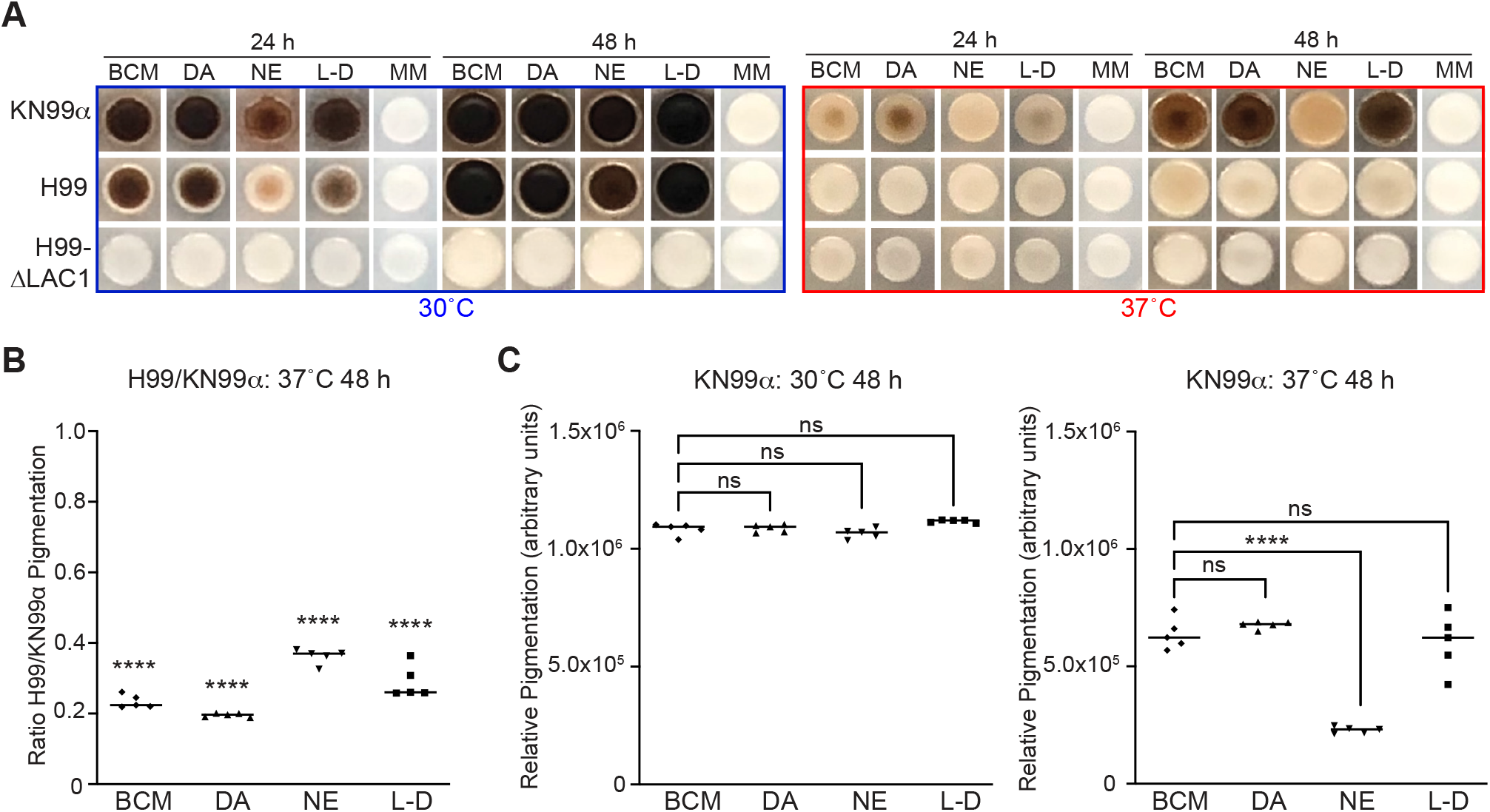
Melanization of *C. neoformans* grown in the presence of different precursors. A, Representative images of *C. neoformans* strains KN99α, H99O, and H99-ΔLAC1 that were spotted on agar supplemented with a mixture of 0.6 mM dopamine, 0.33 mM norepinephrine, and 0.07 mM epinephrine (BCM), 1 mM dopamine (DA), 1 mM norepinephrine (NE), 1 mM L-DOPA (L-D), or minimal media only control (MM) and allowed to melanize at the indicated temperatures for 24 and 48 h. B, Quantified H99O pigmentation expressed as a fraction of that measured for KN99α after 48 h at 37°C showing significantly reduced melanization for H99O compared to KN99α under all conditions tested. C, Quantification of KN99α pigmentation relative to the ΔLAC1 negative control after 48 h at 30°C (left graph) and 37°C (right graph). Statistical significance for this and all subsequent analyses was calculated using an ordinary one-way ANOVA (ns = not significant, *p < 0.05, **p < 0.01, ***p < 0.001, ****p < 0.0001).

### Melanin-containing ‘ghosts’ can be isolated from BCM-supplemented cultures and brain catecholamine extract plates

To further characterize the melanin produced by *C. neoformans* when provided with a mixture of catecholamine precursors, KN99α was melanized in liquid culture for 10 d at 30°C, which is the temperature of optimal growth, in MM supplemented with BCM (0.6 mM DA / 0.33 mM NE / 0.07 mM E) or either 1 mM DA or 1 mM NE. Pigmentation intensity increased more rapidly for cells grown in BCM compared to DA initially but both cultures reached maximal melanization levels after 5 d (Fig. 2A). The culture grown in NE melanized more slowly but eventually reached an intensity comparable to the other two cultures after 10 d (Fig. 2A). Melanization was also achieved for cultures grown in MM supplemented with BCM, DA, or NE at the human body temperature of 37°C, although pigment production occurred more slowly and at a comparable rate for all three cultures (Fig. 2B).

**Figure 2.**
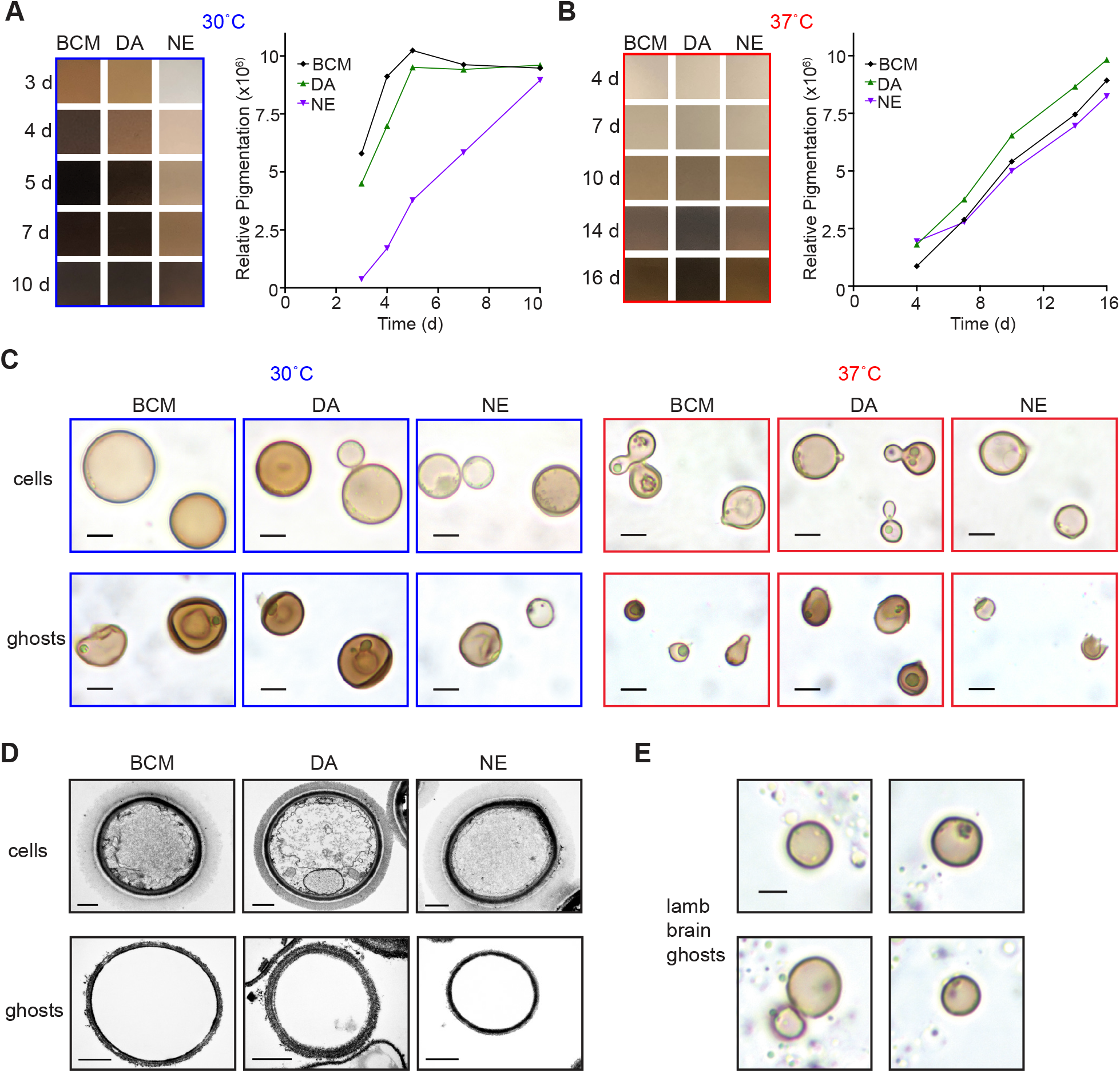
Melanin ghosts from KN99 α melanized in pure or tissue-extracted catecholamines. Color photographs (left) and quantification (right) of relative pigment levels of 2 ml samples taken from KN99α cultures grown for the indicated number of days in minimal media supplemented with a mixture of 0.6 mM dopamine, 0.33 mM norepinephrine, and 0.07 mM epinephrine (BCM), 1 mM dopamine (DA), or 1 mM norepinephrine (NE) at either 30°C (A) or 37°C (B). C, Brightfield images of melanized cells prior to harvesting and acid-resistant melanin ‘ghost’ particles recovered from each culture. D, Transmission electron micrographs of melanized cells and melanin ghost particles from cultures supplemented with the indicated precursors at 30°C. E, Brightfield images of cells melanized on agar plates supplemented with lamb brain catecholamine extract. Scale bars are 5 *μm* and 1 *μm* for light and electron microscopy respectively.

To obtain melanin for biophysical analysis, cells from each culture were harvested and subjected to enzymatic cell wall digestion, protein denaturation and hydrolysis, lipid extraction, and finally boiling in concentrated HCl. In each case, acid-resistant material was recovered that, upon examination by light microscopy, was verified to be dark, yeast-shaped and yeast-sized silhouettes referred to as melanin ‘ghosts’ (Fig. 2C). These particles were also confirmed to be non-viable since plating > 1 × 10^7^ particles on SAB agar yielded no colonies. Microscopic examination revealed differences in the physical appearance of particles derived from each culture. The melanin ghost particles isolated from the DA control culture grown at 30°C were darkly pigmented (Fig. 2C) and similar in appearance to those isolated previously from cultures melanized in L-DOPA at 30°C (39). Fewer intact particles were recovered from the corresponding NE control culture and they tended to be small and pale (Fig. 2C); there was an overall lower yield of acid resistant material (70 mg/L) compared to that of DA (130 mg/L). The BCM melanin ghosts from the culture grown at 30°C were intermediate between the two controls both in appearance, with a mixture of intact dark particles and more fragile, pale particles (Fig. 2C), and in terms of the yield of material (100 mg/L). Ultrastructural examination of melanin ghost particles using transmission electron microscopy revealed ‘empty shells’ with no evidence of membrane-bound organelles or polysaccharide capsule that were visible in the corresponding pre-treatment samples (Fig. 2D).

Melanin ghost particles isolated from cultures melanized at 37°C were smaller and reduced in yield by several-fold (37 mg/L for DA and 14 mg/L for BCM and NE) compared to those isolated at 30°C. The reduced yield is consistent with lower expression of laccase enzyme at 37°C (38). An even smaller yield of intact acid-resistant particles (<1 in 10^6^ of the total number of cells) were recovered from KN99α cells grown at 30°C on agar plates containing brain tissue extract (Fig. 2E). In light of this limitation, the robust pigment production achieved in our large-scale BCM-supplemented culture system, especially at 30°C, supports its utility for acquiring melanin that approximates that produced in human brain tissue but in sufficient quantities to permit structural characterization.

### Solid-state NMR suggests that cells provided BCM incorporate more than one precursor into the melanized cell wall architecture

Given the strong resemblance of the particles derived from *C. neoformans* grown in BCM to melanin ghosts, we sought to characterize the molecular profile of this pigmented material. Solid-state NMR (ssNMR) analysis was implemented to examine its molecular architecture and compare it to that of melanin ghost material isolated from cells grown with either of the two major constituent precursors, DA or NE. The 1D ^13^C cross-polarization magic-angle spinning (CPMAS) spectra of each of these three samples clearly displayed all of the spectral features that are characteristic of melanin ghosts (Fig. 3) (29–31), notably the group of broad, overlapping peaks located between ~110 and 160 ppm. These peaks correspond to the various aromatic carbons that comprise the inherently heterogeneous melanin polymer: although many of the carbons are chemically similar, they differ in both their bonding patterns and local chemical environments, and consequently resonate at slightly different frequencies. This results in a multitude of signals that overlap in the aromatic-carbon spectral region, forming what visually appears as a single, broad peak and serves as a telltale indicator of the presence of melanin pigments. Other characteristic features of the cellular components of melanin ghosts were also observed in all three sample spectra: each of the spectral regions between each ~10-40 ppm and ~50-110 ppm displayed groups of peaks which are attributable to long-chain fatty acids within triacylglycerols and polysaccharide-ring carbons, respectively (40). The exceptional recalcitrance of melanin pigments, which in *C. neoformans* are deposited and intimately interwoven throughout the cell wall, protect this structure from the harsh treatments performed during melanin isolation. Hence, this process yields melanized cell walls (i.e., melanin ‘ghosts’), which contain remnants of the polysaccharide scaffold in addition to trapped neutral lipids, rather than only the purified fungal pigments. Melanin ghost spectra therefore characteristically display ssNMR signals that correspond to non-pigment cellular remnants in addition to those that correspond to the melanin pigment itself. The fact that these signals were observed in all three sample spectra confirmed that the black particles recovered from *C. neoformans* cell cultures grown with BCM are indeed melanin ghosts.

**Figure 3.**
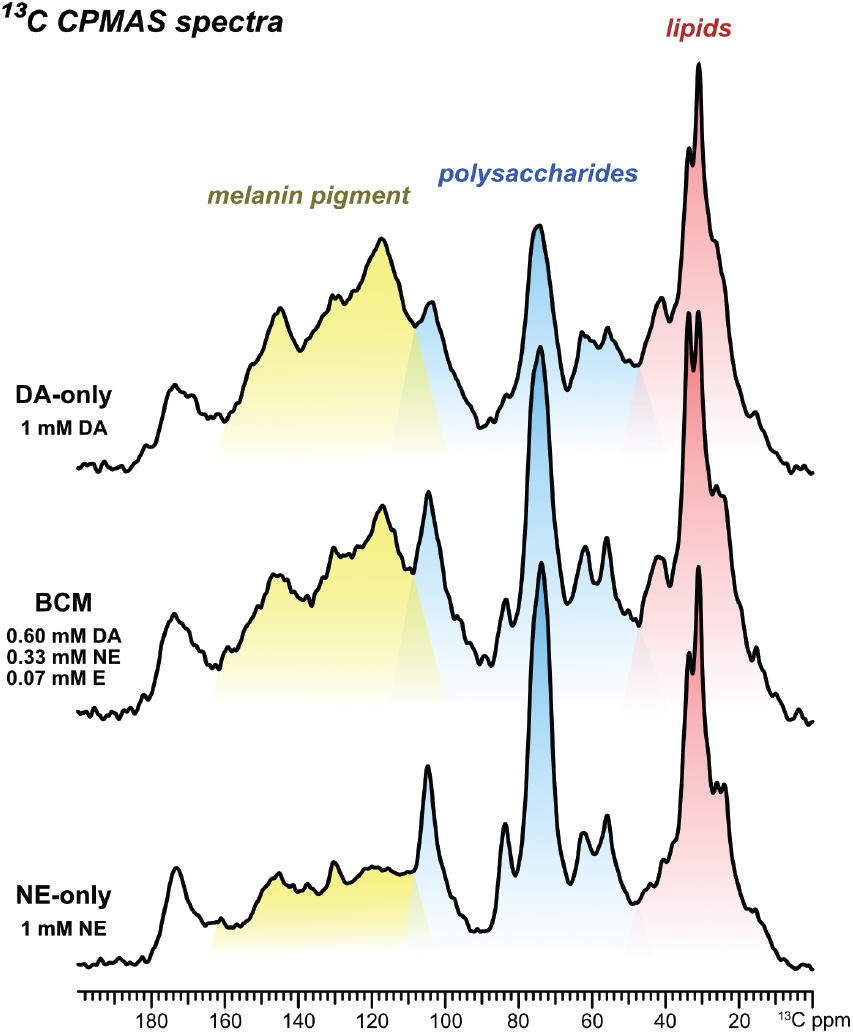
Solid-state NMR spectra of melanin ‘ghosts’ from *C. neoformans* cells grown with DA, NE, or BCM. 1D ^13^C cross-polarization (CPMAS) ssNMR spectra of melanin ghosts isolated from *C. neoformans* KN99α cultures grown for 10 days at 30°C in minimal media containing either 1 mM dopamine (DA, top), 1 mM norepinephrine (NE, bottom) or in a ‘brain catecholamine mixture’ of 0.6 mM dopamine, 0.33 mM norepinephrine, and 0.07 mM epinephrine (BCM, middle). The spectral region shaded in red (10-40 ppm) displays peaks primarily corresponding to the aliphatic carbons of lipids, the region in blue (50-110 ppm) to polysaccharide-ring carbons, and the region in yellow (110-160 ppm) to the aromatic carbons of the melanin pigment. The unshaded region (165-190 ppm) displays several overlapping peaks corresponding to the carbonyl carbons in all three constituent types.

Our ssNMR analysis also provided insight into whether *C. neoformans* is able to incorporate more than one melanization precursor into the pigment architecture when provided with multiple substrates. The heterogenous nature of the melanin polymer which results in multiple broad, overlapping peaks precludes the identification of specific signals that are unique to a melanin synthesized from DA or from NE. Through visual inspection alone it was evident that the DA, NE and BCM melanin ghost samples all share common spectral features but the relative intensities of the groups of peaks corresponding to pigments, lipids and polysaccharides clearly differed. Notably, the aromatic-carbon signals attributed to the pigment contributed substantially less to the total signal intensity in the NE-only melanin ghost spectrum in comparison to the DA-only spectrum, even though both samples were isolated from cell cultures that contained 1 mM concentrations of the respective precursors. This finding was anticipated since prior studies had demonstrated that precursors such as DA or L-DOPA result in more robust melanin production in comparison to others such as NE or E (29–31). This, in turn, results in melanin ghosts that have a greater pigment content relative to the other cellular remnants present such as polysaccharide and lipids. The spectrum of the BCM melanin ghosts, which were isolated from a cell culture containing 0.6 mM DA, 0.33 mM NE and 0.07 mM E, exhibited pigment contributions with respect to other cellular constituents that exceeded expectations based on utilization of only the robust DA precursor, suggesting incorporation of the additional catecholamines. Quantitatively reliable direct polarization (DPMAS) spectra reinforced the trend of higher-than-expected pigment contributions in the melanin ghosts (Fig. S1). Moreover, this trend was also observed in both the CPMAS and DPMAS spectra of melanin ghost samples isolated from *C. neoformans* cells grown at 37°C despite the overall reduction in melanin production that occurs at this temperature (Fig. S2). Thus, even without detailed discrimination of the pigment structures synthesized from different precursors, our ssNMR analysis suggested that *C. neoformans* can use at least two different melanization precursors when they are provided simultaneously.

### Radiolabeled norepinephrine and dopamine are incorporated into C. neoformans melanin

As our structural data provided highly suggestive evidence for the incorporation of multiple precursors, we extended our analysis by employing radiolabeled catecholamines as a sensitive and quantitative means of determining whether both DA and NE can be incorporated into the melanin pigment at the same time. Individual *C. neoformans* KN99α cultures were grown in the presence of a low concentration (0.0001 mM) of high specific activity ^3^H-labeled NE with and without the addition of unlabeled NE or DA. A control culture deficient in the primary laccase enzyme (ΔLAC1) was also supplemented with the same concentration of ^3^H-NE as a negative control. Even though the molar concentration of NE in the culture was very low, we detected significantly more radioactivity retained in the cell pellet from the KN99α cell culture grown without any additional unlabeled precursors compared to that of the ΔLAC1 control culture (Fig. 4A, graph). The addition of excess unlabeled NE at a concentration of either 0.1 or 1 mM resulted in a noticeable, concentration-dependent increase in pigmentation (Fig. 4A, lower panels) and a concomitant increase in the incorporation efficiency of ^3^H-NE (Fig. 4A, graph). We attribute these increases to the established dose dependence of melanization for DA and NE precursors, possibly resulting from induction of the laccase enzyme (41). Notably, a comparable increase in ^3^H-NE incorporation was also detected when cultures were supplemented with excess (1 mM) of unlabeled DA (Fig. 4A, graph), indicating that *C. neoformans* can utilize more than one type of precursor simultaneously.

**Figure 4.**
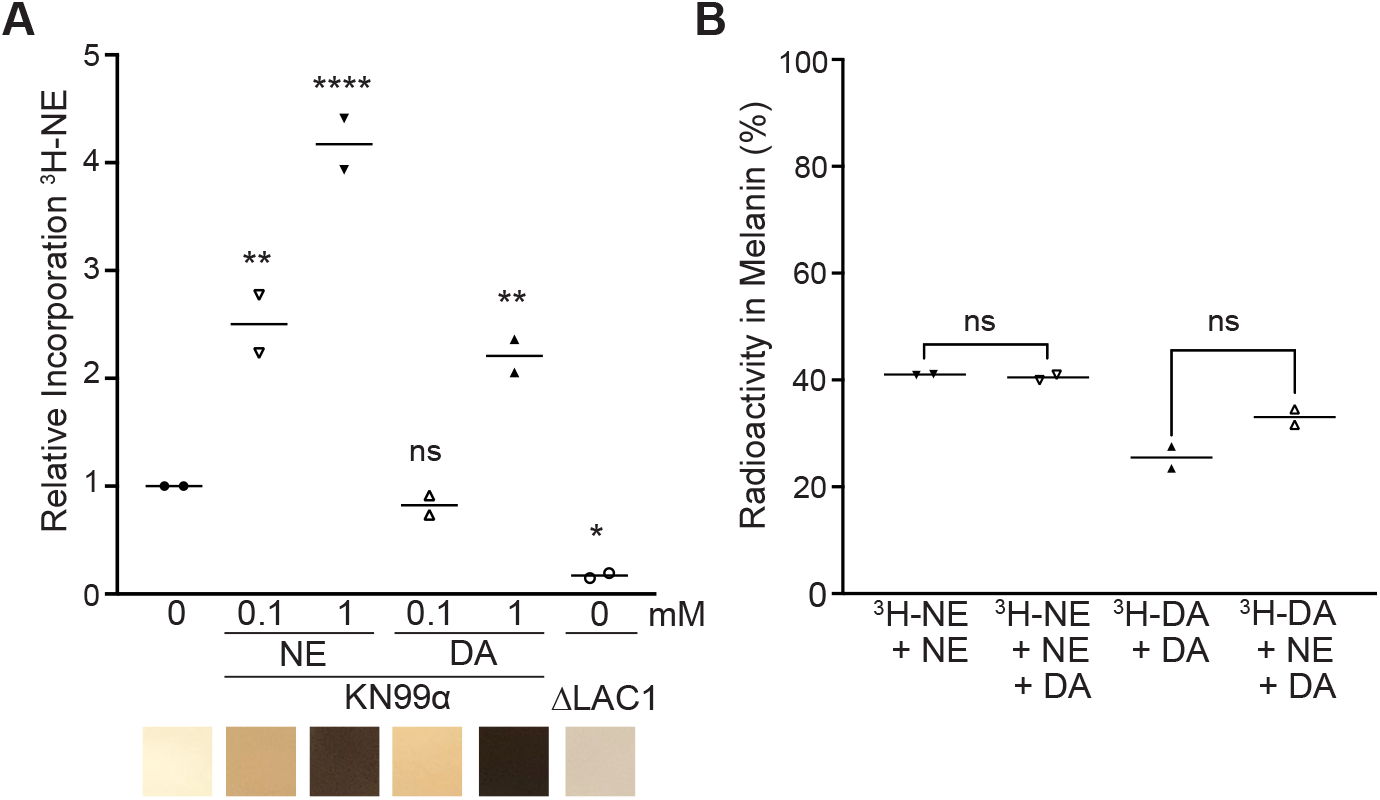
Incorporation of radiolabeled precursors into melanin. A, Scatter plot graph (upper panel) of the amount of radioactivity retained in cell pellets of KN99α and ΔLAC1 cultures supplemented with 2 *μ*Ci ^3^H-NE (0.0001 mM) in the absence or presence of the indicated concentration of unlabeled catecholamines as expressed relative to cultures with ^3^H-NE only (n = 2). Panel below graph displays representative regions taken from color photographs of each culture. B, Radioactivity measured in acid-resistant melanin ghost particles as a percentage of the radioactivity in the samples before acid treatment. Cultures were melanized in minimal media supplemented with 0.0001 mM ^3^H-NE or ^3^H-DA and with 1 mM unlabeled catecholamines as indicated (n = 2).

To confirm that the radioactivity associated with *C. neoformans* cell pellets resulted from the incorporation of the radiolabeled precursors into melanin, we heated cells that had been melanized in mixtures of radiolabeled and unlabeled precursors in concentrated hydrochloric acid to generate melanin ghost particles. Approximately 30-40% of the radioactivity was retained in melanin from cells labeled with either ^3^H-NE or ^3^H-DA (Fig. 4B). The efficiency of incorporation of either radioactively-labeled precursor into melanin was not significantly different when melanization was supported by an excess of a single unlabeled catecholamine or a 1:1 mixture of NE and DA (Fig. 4B) since in either case, a substantial proportion of the ^3^H radioactivity was retained after HCl treatment.

### Melanin derived from different precursors provide similar protection from UV radiation and oxidative damage

Melanization of *C. neoformans* in L-DOPA was previously demonstrated to afford protection from the fungicidal effects of UV radiation (42). To investigate the protective capacity of the brain mixture of catecholamines, KN99α cells were melanized for 7 d in BCM, DA, NE, or L-DOPA and compared to non-melanized control cells for survival after UV exposure. Cells melanized under all conditions showed significantly higher percentages of survival compared to non-melanized cells, although melanization in NE was less protective than with the other three precursors (Fig. 5A). Melanization in BCM showed no statistical difference from DA or L-DOPA in shielding cells from UV radiation (P values of 0.8107 and 0.6413 respectively).

**Figure 5.**
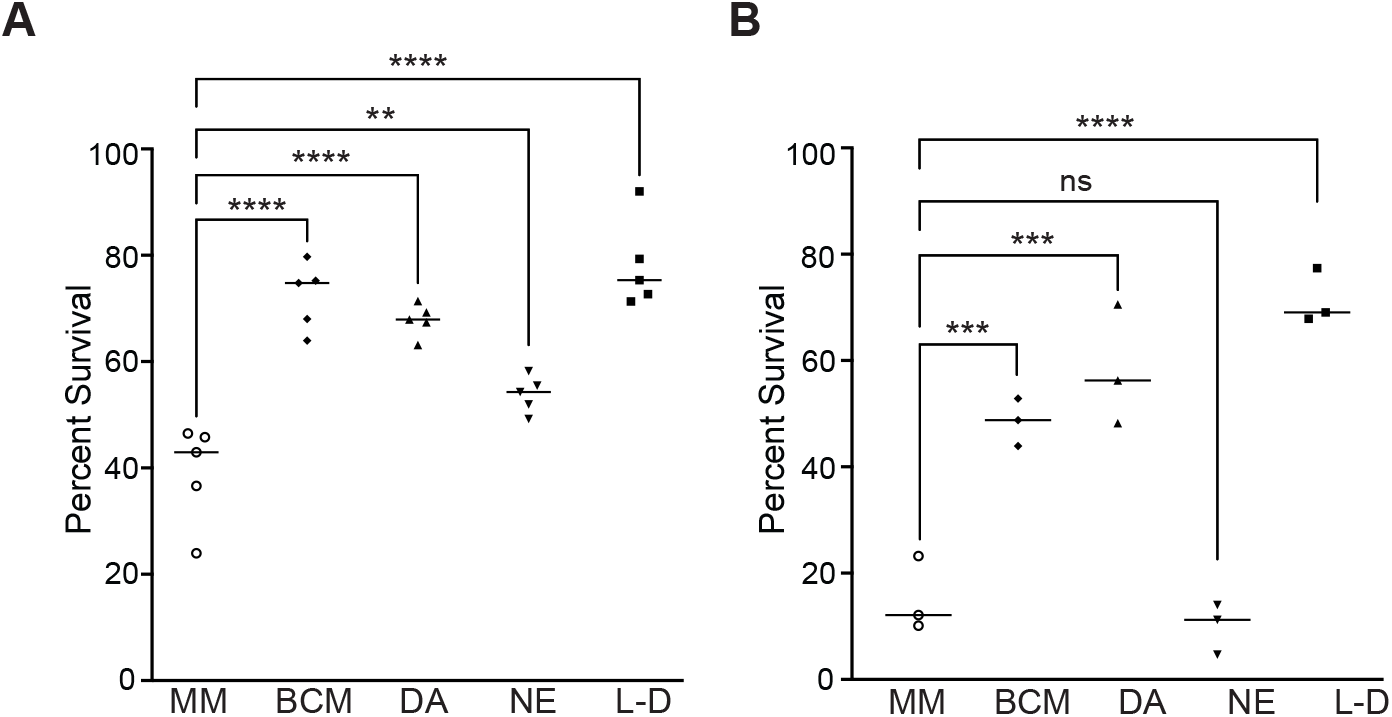
Comparison of melanization conditions for resistance to cellular damage. A, Percent survival relative to untreated controls for melanized and non-melanized (MM) KN99α plated on SAB agar plates and then exposed to 15,000 *μ*J/cm^2^ UV radiation (n = 5). B, Survival of KN99α melanized with the indicated catecholamines or a non-melanized control (MM) was determined as the number colony forming units (CFU) for cultures exposed to oxygen-derived radicals for 5 hours at 37°C as a percentage of the CFU for untreated controls (n = 3). Melanization conditions were 0.6 mM dopamine, 0.33 mM norepinephrine, and 0.07 mM epinephrine (BCM), 1 mM dopamine (DA), 1 mM norepinephrine (NE), and 1 mM L-DOPA (L-D).

Since melanin is a key virulence factor that contributes to *C. neoformans* survival inside human host cells, we sought to explore the ability of ‘natural’ melanin to safeguard against damaging free radicals generated inside macrophages. An *in vitro* oxidative system that produces oxygen-derived free radicals (43) was used to treat KN99α cells that were non-melanized or had been melanized for 7 d in the BCM, DA, NE, or L-DOPA. Melanization in the presence of each of these precursors, with the exception of NE, significantly increased the percentage of cells that survived exposure to oxidative damage at 37°C compared to non-melanized cells (Fig. 5B).

## Discussion

Melanization by *C. neoformans* is a prominent virulence factor that facilitates evasion from the host immune system. Melanin produced by *C. neoformans* grown in defined culture media supplemented with the phenolic precursor, L-DOPA, has been well characterized, but much less is known about the protective layer of pigment deposited during infection. In the brain, L-DOPA is readily converted to dopamine, which is in turn a precursor of norepinephrine and epinephrine. Hence, *C. neoformans* cells that disseminate to brain tissue are expected to utilize one or more of these three catecholamines for melanin synthesis. We cultured *C. neoformans* in a mixture of dopamine, norepinephrine and epinephrine reflective of their relative proportions in the human brain and found that the resulting melanin was similar to L-DOPA melanin in its ability to protect cells from UV radiation and oxidative damage. Structural analysis using solid-state NMR revealed a spectrum consistent with the simultaneous incorporation of more than one precursor. Melanization experiments employing radiolabeled catecholamines provided additional evidence that *C. neoformans* can use multiple precursors simultaneously and suggested that melanin made during infection is likely synthesized from a mixture of the available substrates.

Initial evidence for *C. neoformans* melanization during infection was provided by the isolation of acid-resistant melanin ghost particles from infected animal and human tissue and from cells grown on tissue homogenate agar plates (28, 33). We isolated similar particles from cells grown on agar plates infused with a lamb brain catecholamine extract but only a very small fraction of the total cells was recovered as melanin ghosts. Neurotransmitters such as dopamine in brain tissue are contained within vesicles and degraded by monoamine oxidase, an enzyme found in glial cells (44). Consequently, several factors could have contributed to the small yield when using brain tissue as the source of substrate, including the relatively lower concentrations of catecholamines in brain tissue compared to the concentrations achievable in culture, low bioavailability of the precursors sequestered in neurotransmitter vesicles, and degradation of catecholamines during the extraction procedure when cells are lysed. In light of these limitations, we opted for culturing *C. neoformans* in a mixture of catecholamines proportional to their relative concentrations present in the human brain and achieved robust melanization on both solid agar substrates and in liquid cultures. We also observed more efficient melanization by the KN99α variant compared to its parental H99O strain, especially at the human body temperature of 37°C, and this is consistent with the correlation between melanization and virulence (36).

Although *C. neoformans* is able to produce melanin pigments from a variety of aromatic substrates, certain precursors, e.g., L-DOPA and dopamine, promote more robust melanization compared to norepinephrine or epinephrine, as evidenced by a greater yield by mass melanin ghosts that have relatively thicker “shells” and a larger stable free-radical population (39). In our prior work, ssNMR analysis was implemented to determine whether *C. neoformans* is able to distinguish between different melanization precursors when two are provided at the same time (31). We found that *C. neoformans* will selectively use a more favorable precursor (L-DOPA) over a less favorable one (norepinephrine) when each is provided at a 1 mM concentration. Although we had determined that *C. neoformans* will preferentially utilize the more favorable L-DOPA precursor, we did not determine whether this occurred with the exclusion of norepinephrine utilization, or if the outcome would differ if the total concentration of both precursors was 1 mM. In our current work, we used ssNMR to examine the melanin ghosts isolated from cells grown in a mixture of catecholamines in the proportions found in the human brain (60% dopamine, 33% norepinephrine, and 7% epinephrine (34, 35)) and provided at a total precursor concentration of 1 mM. As observed for ghosts isolated from cultures containing 1 mM of either dopamine or norepinephrine, the ^13^C CPMAS spectrum of these ‘brain catecholamine mix’ (BCM) ghosts displayed all the hallmark features of *C. neoformans* melanized cell walls, including signals that correspond to pigments, polysaccharides, and lipids (Fig. 3). However, the relative intensities of the signals attributed to each of these constituents, which are indicative of their approximate relative quantities, differed among the three ghost spectra. The analysis of quantitatively-reliable ^13^C DPMAS spectra verified this observation. As anticipated, the ghosts prepared from the culture containing 1 mM of the less-favorable norepinephrine precursor had the lowest relative pigment content whereas those prepared from the 1 mM dopamine culture had the highest content. Remarkably, the ghosts isolated from the BCM culture, which only contained 0.6 mM dopamine, were found to have a pigment content nearly equivalent to that of ghosts derived from a culture provided with 1 mM dopamine, and thus suggested that one or both of the other precursors (norepinephrine and epinephrine) present in the mixture were being utilized for pigment formation in addition to dopamine.

Our interpretation of the ssNMR quantitative analysis assumes that 100% of the melanization precursors provided in culture media are utilized for melanin formation. But if the complement of provided precursors influences the efficiency with which a particular precursor is utilized, our results could also be interpreted to suggest increased relative dopamine incorporation for cells provided with the brain catecholamine mixture compared to dopamine alone. To address more directly whether *C. neoformans* can incorporate more than one catecholamine simultaneously, we cultured cells in the presence of radiolabeled catecholamines to follow their incorporation into melanin. A significantly higher level of radioactivity was detected in the cell pellet of the wild-type KN99α strain compared to a laccase-deficient mutant, ΔLAC1, which suggests that the labeled precursor was used to support melanization. However, the efficiency of incorporation by KN99α was very low at approximately 5%, which is not unexpected given the low molar concentration of 0.0001 mM for the labeled precursor. *C. neoformans* melanization in culture was previously shown to have a dose-dependence for L-DOPA and norepinephrine substrates, with an optimal concentration of 1 mM (41). In this instance, supplementing cultures of KN99α with 0.1 and 1 mM unlabeled norepinephrine in combination with 0.0001 mM ^3^H-labeled norepinephrine resulted in a concentration-dependent increase in both visible pigmentation and incorporation of ^3^H-NE. Remarkably, melanization with 1 mM unlabeled DA also significantly increased the incorporation of ^3^H-labeled norepinephrine, although not to the same extent as 1 mM norepinephrine, possibly reflecting a slight incorporation bias for dopamine. However, the increased efficiency of ^3^H-labeled norepinephrine incorporation in the presence of a 10,000-fold molar excess of dopamine supports the conclusion that *C. neoformans* does not use dopamine to the exclusion of norepinephrine. To confirm that the radiolabeled precursors were being incorporated into melanin pigment and not just peripherally associated with cells, we isolated acid-resistant melanin ghosts from radiolabeled cultures. Using either ^3^H-labeled norepinephrine or ^3^H-labeled dopamine, we were able to isolate radiolabeled melanin when cells were melanized in either individual unlabeled catecholamines or an equimolar mixture of the two precursors. Together, these results suggest that the most readily abundant substrate is used to establish a melanin scaffold that is then built upon using any and all available precursors, thereby generating local structural diversity for the melanin particles.

The study of *C. neoformans* melanization has used predominantly L-DOPA as a substrate, but this precursor would not be the major substrate available in host tissues. *C. neoformans* infections are initially established in the lungs where they encounter alveolar macrophages. As these and other immune cells are known to produce and release catecholamines (45), it is possible that melanization supported by lung resident catecholamines could enhance virulence and promote brain dissemination. The ability of *C. neoformans* to incorporate multiple catecholamines simultaneously suggests that melanin synthesized in brain tissue may differ in distinct anatomical regions as the relative proportions of these neurotransmitters have been found to vary widely from one area of the brain to another (34, 35). Intriguingly, *C. neoformans* infections are often found to be concentrated in the basal ganglia region of the brain where dopamine levels are highest (46, 47), a finding that could reflect resistance of melanized cells to clearance by host immune mechanisms. Basal ganglia preference is also consistent with the notion that melanin produced by *C. neoformans* in the brain is primarily dopamine-derived. Consequently, we suggest that future studies of the role of melanin in *C. neoformans* pathogenesis are more likely to reflect brain infection conditions if dopamine is used as the melanization substrate. However, L-DOPA may be a more suitable substrate to study the role of cryptococcal melanization in the environment where L-DOPA is a common precursor of plant compounds and accumulates in soils (48). We note that although the type of melanin produced by *C. neoformans* is dependent on the available precursor, our prior ssNMR studies revealed considerable structural similarity among different melanins (29–31).

An improved understanding of the composition and structure of ‘natural’ melanin produced by *C. neoformans* during infection will help elucidate its function in virulence. We have shown that *C. neoformans* can incorporate multiple precursors, even when their molar ratios differ by several orders of magnitude. Given that melanin synthesis is driven by the accretion of highly reactive intermediates into a solid material, it is expected that *C. neoformans* provided with a mixture of precursors produces polytypic melanin. Furthermore, depending on the relative availability of specific catecholamines, structurally diverse forms of melanin may be produced in different anatomical regions during the course of infection. Despite this added level of complexity, melanins produced in culture from single or mixed substrates were found to be functionally comparable in their capacity to protect *C. neoformans* from UV radiation and oxidative damage, but whether specific forms of melanin could imbue *C. neoformans* with a greater survival advantage during infection remains unanswered. Whereas the present study has shed light on initial polymerization events, the resulting melanin compound is a highly complex, amorphous substance for which detailed structural elucidation remains beyond our analytical horizon.

## Experimental Procedures

### Yeast strains

*Cryptococcus neoformans* serotype A strains H99O (American Type Culture Collection 208821), H99-ΔLAC1 (Fungal Genetics Stock Center, 2007 Lodge library), and KN99α (Fungal Genetics Stock Center, 2007 Lodge library) were recovered from frozen 50% glycerol stocks by growth in YPD broth (Difco 242820) for 48 h at 30°C. The hygromycin resistance cassette in H99-ΔLAC1 was retained by supplementing culture media with 200 U/ml Hygromycin B from *Streptomyces sp.* (Calbiochem 400051). Cells were washed twice with sterile phosphate-buffered saline before sub-culturing into defined media.

### Culture media with relative proportions of catecholamines in the human brain

To study melanization under conditions representative of the human brain, we used the total cortical and subcortical measurements of the three brain catecholamines reported previously (34, 35) to calculate the relative proportions of approximately 60% dopamine (DA), 33% norepinephrine (NE), and 7% epinephrine (E). *C. neoformans* variant KN99α was grown in minimal media (MM) consisting of 29.4 mM K_2_HPO_4_, 10 mM MgSO_4_, 13 mM glycine, 15 mM D-glucose, and 3 μM thiamine at pH 5.5 that was supplemented with the brain catecholamine mixture (BCM) consisting of 0.6 mM Dopamine hydrochloride (MilliporeSigma H8502), 0.33 mM (-)-Norepinephrine (MilliporeSigma A7257), and 0.07 mM (-)-Epinephrine (MilliporeSigma E4250). Control cultures were also grown in MM supplemented with 1.0 mM DA or 1.0 mM NE.

### Melanization assay on agar plates

To prepare melanization assay plates, 4X MM with no additives or with either 1.0 mM DA, 1.0 mM NE, 1.0 mM 3,4-Dihydroxy-L-phenylalanine (L-DOPA, MilliporeSigma D9628), or BCM was mixed 1:3 with cooled 2% agar (Difco 214530) and poured into individual wells of 24-well plates (2 ml per well). Plates were allowed to cool at RT for 5 h, then 1 × 10^6^ PBS-washed KN99α, H99O, or H99-ΔLAC1 cells were pipetted into each well and plates were incubated at either 30°C or 37°C. After 24 and 48 h, the plates that had been incubated at each temperature were photographed in a single field using a 12-megapixel camera. Images were converted to grayscale in Adobe Photoshop (version 22.0.1) and intensities were quantified using Image Studio Lite software (version 5.2.5, Li-Cor Biosciences). The signal measured for the MM control was applied as the image background and intensity measurements for the H99-ΔLAC1 negative control were subtracted from those of KN99α and H99O to derive pigmentation levels above background.

### Isolation of melanin particles

Cultures of KN99α were grown in MM supplemented with either BCM, 1.0 mM DA, or 1.0 mM NE at 30°C or 37°C. Volumes of 0.5 L for BCM and DA and 1 L for NE for cultures grown at 30°C were doubled for growth at 37°C. Melanization of the cultures was monitored by photographing and quantifying pigment intensities (as described above) for 2 mL samples that had been transferred to the wells of a transparent 12-well plate (Celltreat, 229112). After incubation for 10 d at 30°C or 16 d at 37°C, acid-resistant melanin ‘ghost’ particles were isolated from cultures as described in detail previously (29, 33). Briefly, cells were treated with a 10 mg/ml solution of *Trichoderma harzianum* lysing enzymes (MilliporeSigma L1412) in 1.0 M sorbitol and 0.1 M sodium citrate for 24 h at 30°C to remove the fungal cell wall. The resulting protoplasts were subjected to protein denaturation with 4.0 M guanidinium thiocyanate for 18 h at RT and then proteolysis with a 20 mg/ml solution of proteinase K (MilliporeSigma 03115801001) in 10 mM Tris-HCl (pH 8), 5.0 mM CaCl_2_, and 0.5% sodium dodecyl sulfate for 4 h at 65°C. Following lipid removal by three successive extractions with a 4:8:3 ratio of methanol-chloroform-saline (49) samples were heated at 90°C in 6 M HCl for 1 h. Melanin-containing particles were recovered by centrifugation and dialyzed for several days against pure H_2_O prior to lyophilization. Dry material was weighed using an analytical balance (Mettler Toledo AT261 DeltaRange) to determine yield.

### Melanization in brain tissue extracts

The tissue from three lamb brains (227 g) was homogenized in ~1 ml/g ice cold buffer comprised of 0.1 M perchloric acid, 1.0 mM sodium metabisulfite, and 0.1 mM EDTA using a 30 ml Potter-Elvehjem tissue grinder (Wheaton 358011) attached to a drill press. Following centrifugation at 5000 rpm for 20 min at 4°C, the pH of the supernatant was adjusted to ~5 by dropwise addition of sodium hydroxide and then passed through a 0.22 μm filter (Millipore SCGPS02RE). The filtrate was rapidly frozen on dry ice and then lyophilized for ~72 h. The freeze-dried extract was resuspended with 75 ml 2X MM containing 1% Penicillin-Streptomycin, mixed with 75 ml cooled 3% agar and poured into six 100 ml petri dishes. After ~24 h at RT, plates were spotted with PBS-washed KN99α and incubated at 30°C for 26 d. Melanin ghost particles were isolated from cells melanized in liquid cultures and on plates as described previously.

### Light and transmission electron microscopy

Melanized cells and melanin ghost particles were imaged under bright field illumination with an Olympus AX70 microscope (Olympus, Center Valley, PA) using an oil immersion 100X objective. Images were captured using QCapture-Pro 6.0 software with a Retiga 1300 digital CCD camera. (Teledyne Photometrics, Tucson, AZ). For transmission electron microscopy (TEM), samples were fixed in a buffer comprised of 2.5% glutaraldehyde, 3 mM MgCl_2_ and 0.1 M sodium cacodylate (pH 7.2) overnight at 4°C. After buffer rinse, samples were postfixed in 1% osmium tetroxide, 1.25% potassium ferrocyanide in 0.1 M sodium cacodylate for at least one hour (no more than two) on ice in the dark. After the fixing step, samples were rinsed in dH_2_O, followed by uranyl acetate (2%, aq.) (0.22 μm filtered, 2.5 hr, dark), dehydrated in a graded series of ethanol and embedded in Spurrs (Electron Microscopy Sciences) resin. Samples were polymerized at 60°C overnight. Thin sections, 60 to 90 nm, were cut with a diamond knife on a Leica UCT ultramicrotome and picked up with 2 × 1 mm Formvar copper slot grids. Grids were stained with 2% aqueous uranyl acetate followed by lead citrate and observed with a Hitachi 7600 TEM at 80 kV. Images were captured with an AMT CCD XR80 (8 megapixel side mount AMT XR80 high-resolution high-speed camera).

### Solid-state NMR of melanin particles

Solid-state NMR measurements were carried out on a Varian (Agilent) DirectDrive2 (DD2) spectrometer operating at a ^1^H frequency of 600 MHz. All data were acquired on ~8-10 mg of lyophilized material using a 1.6-mm T3 HXY fastMAS probe (Agilent Technologies, Santa Clara, CA) with a magic-angle spinning (MAS) rate of 15.00 ± 0.02 kHz and a spectrometer-set temperature of 25 °C. The 1D ^13^C cross-polarization (CPMAS) experiments were conducted with 90° pulse lengths of 1.5 and 1.7 μs for ^1^H and ^13^C, respectively, a 10% linearly ramped amplitude for ^1^H and a 1 ms contact time for ^1^H-^13^C polarization transfer. The 1D ^13^C direct polarization (DPMAS) experiments were conducted with 90° pulse lengths of 1.2 and 1.4 μs for ^1^H and ^13^C, respectively, and long recycle delays (50 s) were used in order to obtain quantitatively reliable signal intensities. Heteronuclear decoupling was applied during signal acquisition using the small phase incremental alternation pulse sequence (SPINAL) (50) and a radio-frequency field strength of 104 kHz.

### Radiolabeled norepinephrine incorporation analysis

KN99α was grown in MM for 2 d at 30°C, after which cells were washed and resuspended in fresh media at a cell density of 2 × 10^8^ cells/ml. Cultures were labeled with 2 μCi/ml (0.0001 mM) norepinephrine DL-[7-^3^H] hydrochloride (^3^H-NE, American Radiolabeled Chemicals, Inc. ART-0172) for 7 d at 30°C in the absence or presence of 0.1 or 1.0 mM unlabeled NE or 0.1 or 1.0 mM unlabeled DA. H99-ΔLAC1 was also labeled with ^3^H-NE as a negative control. Color photographs were taken of each culture to document relative pigmentation intensities. Labeled cells were collected by centrifugation at 16,000 × g for 5 min, washed twice with MM, then resuspended in MM. Samples of media, wash buffer, and resuspended pellets were mixed with UniverSol-ES liquid scintillation cocktail (MP Biomedicals 01882480) and decays per minute (DPM) were quantified using an LS 6500 liquid scintillation counter (Beckman, Indianapolis, IN). Percent incorporation was calculated by dividing the DPM measured for the pellet sample by the total DPM of the pellet, media, and wash samples.

### Isolation of radiolabeled melanin particles

KN99α was grown as described above in MM supplemented with 2 μCi/ml ^3^H-NE and 1.0 mM unlabeled NE or a mixture of 1.0 mM unlabeled NE and DA or with 4 μCi/mL (0.0001 mM) Dihydroxyphenylethylamine 3,4-[Ring-2,5,6-^3^H] (^3^H-DA, Perkin Elmer NET673250UC) and 1.0 mM unlabeled DA or 1.0 mM unlabeled NE and DA for 10 d at 30°C. Labeled cells were collected by centrifugation at 16,000 × g for 10 min then treated with 6 M hydrochloric acid in a 90°C water bath for 1 h. Acid resistant melanin ghost material was collected by centrifugation at 16,000 × g for 10 min. DPM were measured as described above for samples taken before and after acid treatment and the percent radioactivity retained in the melanin ghost particles compared to pre-treatment samples was calculated.

### Exposure to ultraviolet radiation

Equal numbers of non-melanized and 7-d melanized KN99α were plated on Sabouraud dextrose agar (Difco 210950) and then exposed to 15,000 μJ/cm^2^ UV radiation (254 nm) using a Stratalinker 1800 (Stratagene, La Jolla, CA). Percent survival was calculated from CFU counts of treated compared to untreated control plates after incubation at 30°C for 3 d.

### Exposure to oxygen-derived oxidants

Cultures of KN99α were grown at 30°C for 7 d in MM (control) or MM supplemented with BCM, 1.0 mM DA, 1.0 mM NE, or 1.0 mM L-DOPA. After washing with PBS, 2.5 × 10^4^ cells were treated for 5 h at 37°C with an oxidative system that uses epinephrine as an electron donor, H_2_O_2_ as an electron acceptor, and Fe^3+^ as a transition metal catalyst (43); reaction among these three components results in the formation of Fe(III)-epinephrine complexes that are a primary source of oxidative damage in biological systems and have been linked to various stress-related pathologies in mammals (51). A fresh stock solution of 5.0 mM Fe^3+^ was prepared by mixing 10 mM ferric ammonium sulfate, NH_4_Fe(SO_4_)_2_, 1:1 with 1.0 mM sulfuric acid. The oxidative treatment solution consisted of 0.5 mM Fe^3+^, 1.0 mM H_2_O_2_, and 1.0 mM epinephrine bitartrate (MilliporeSigma E4375) in a buffer comprised of 50 mM sodium acetate (pH 5.5) and 1.0 mM MgSO_4_. Percent survival was calculated from counts of colony-forming units (CFUs) for each condition compared to untreated controls.

### Statistical Analysis

Biological replicates were performed for experiments as indicated and statistical significance was determined using an ordinary one-way analysis of variance (ANOVA) with Sidak’s multiple comparisons test (GraphPad Prism 9 software).

## Data Availability

All data are contained within the article.

## Acknowledgments

We are grateful to the Hardwick lab for providing laboratory space for radioisotope experiments and to the Jacobs-Lorena lab for providing us with a UV Stratalinker 1800. We thank Barbara Smith of the Johns Hopkins Microscope Core Facility for providing expert assistance with TEM imaging. This work was supported by the National Institutes of Health, grant numbers R01-AI052733 (A.C and R.E.S.). C.C. was also supported by the Brescia Fund of the CCNY Department of Chemistry and Biochemistry.

## REFERENCES

1. Springer, D. J., Mohan, R., and Heitman, J. (2017) Plants promote mating and dispersal of the human pathogenic fungus Cryptococcus. PLoS One. 12, e0171695

2. Lin, X., and Heitman, J. (2006) The biology of the Cryptococcus neoformans species complex. Annu. Rev. Microbiol. 60, 69–105

3. Coelho, C., Bocca, A. L., and Casadevall, A. (2014) The intracellular life of Cryptococcus neoformans. Annu. Rev. Pathol. 9, 219–238

4. Rajasingham, R., Smith, R. M., Park, B. J., Jarvis, J. N., Govender, N. P., Chiller, T. M., Denning, D. W., Loyse, A., and Boulware, D. R. (2017) Global burden of disease of HIV-associated cryptococcal meningitis: an updated analysis. Lancet. Infect. Dis. 17, 873–881

5. May, R. C., Stone, N. R. H., Wiesner, D. L., Bicanic, T., and Nielsen, K. (2016) Cryptococcus: from environmental saprophyte to global pathogen. Nat. Rev. Microbiol. 14, 106–117

6. Chrétien, F., Lortholary, O., Kansau, I., Neuville, S., Gray, F., and Dromer, F. (2002) Pathogenesis of cerebral Cryptococcus neoformans infection after fungemia. J. Infect. Dis. 186, 522–530

7. Chang, Y. C., Stins, M. F., McCaffery, M. J., Miller, G. F., Pare, D. R., Dam, T., Paul-Satyaseela, M., Kim, K. S., and Kwon-Chung, K. J. (2004) Cryptococcal yeast cells invade the central nervous system via transcellular penetration of the blood-brain barrier. Infect. Immun. 72, 4985–4995

8. Perfect, J. R., Dismukes, W. E., Dromer, F., Goldman, D. L., Graybill, J. R., Hamill, R. J., Harrison, T. S., Larsen, R. A., Lortholary, O., Nguyen, M.-H., Pappas, P. G., Powderly, W. G., Singh, N., Sobel, J. D., and Sorrell, T. C. (2010) Clinical Practice Guidelines for the Management of Cryptococcal Disease: 2010 Update by the Infectious Diseases Society of America. Clin. Infect. Dis. 50, 291–322

9. Datta, K., and Pirofski, L. (2006) Towards a vaccine for Cryptococcus neoformans: principles and caveats. FEMS Yeast Res. 6, 525–536

10. Iyer, K. R., Revie, N. M., Fu, C., Robbins, N., and Cowen, L. E. (2021) Treatment strategies for cryptococcal infection: challenges, advances and future outlook. Nat. Rev. Microbiol. 19, 454–466

11. Eisenman, H. C., and Casadevall, A. (2012) Synthesis and assembly of fungal melanin. Appl. Microbiol. Biotechnol. 93, 931–940

12. Hill, H. Z. (1992) The function of melanin or six blind people examine an elephant. Bioessays. 14, 49–56

13. Casadevall, A., Rosas, A. L., and Nosanchuk, J. D. (2000) Melanin and virulence in Cryptococcus neoformans. Curr. Opin. Microbiol. 3, 354–358

14. Nosanchuk, J. D., and Casadevall, A. (2006) Impact of melanin on microbial virulence and clinical resistance to antimicrobial compounds. Antimicrob. Agents Chemother. 50, 3519–3528

15. Eisenman, H. C., Frases, S., Nicola, A. M., Rodrigues, M. L., and Casadevall, A. (2009) Vesicle-associated melanization in Cryptococcus neoformans. Microbiology. 155, 3860–3867

16. Eisenman, H. C., Nosanchuk, J. D., Webber, J. B. W., Emerson, R. J., Camesano, T. A., and Casadevall, A. (2005) Microstructure of cell wall-associated melanin in the human pathogenic fungus Cryptococcus neoformans. Biochemistry. 44, 3683–3693

17. Cordero, R. J. B., and Casadevall, A. (2017) Functions of fungal melanin beyond virulence. Fungal Biol. Rev. 31, 99–112

18. Fu, M. S., Liporagi-Lopes, L. C., dos Santos Júnior, S. R., Tenor, J. L., Perfect, J. R., Cuomo, C. A., and Casadevall, A. (2020) Amoeba predation of *Cryptococcus neoformans* results in pleiotropic changes to traits associated with virulence. bioRxiv. 10.1101/2020.08.07.241190

19. Wang, Y., Aisen, P., and Casadevall, A. (1995) Cryptococcus neoformans melanin and virulence: mechanism of action. Infect. Immun. 63, 3131–3136

20. Blasi, E., Barluzzi, R., Mazzolla, R., Tancini, B., Saleppico, S., Puliti, M., Pitzurra, L., and Bistoni, F. (1995) Role of nitric oxide and melanogenesis in the accomplishment of anticryptococcal activity by the BV-2 microglial cell line. J. Neuroimmunol. 58, 111–116

21. Liu, L., Tewari, R. P., and Williamson, P. R. (1999) Laccase protects Cryptococcus neoformans from antifungal activity of alveolar macrophages. Infect. Immun. 67, 6034–6039

22. Polacheck, I., Hearing, V. J., and Kwon-Chung, K. J. (1982) Biochemical studies of phenoloxidase and utilization of catecholamines in Cryptococcus neoformans. J. Bacteriol. 150, 1212–1220

23. Lee, S. C., Dickson, D. W., and Casadevall, A. (1996) Pathology of cryptococcal meningoencephalitis: analysis of 27 patients with pathogenetic implications. Hum. Pathol. 27, 839–847

24. Coelho, C., and Casadevall, A. (2016) Cryptococcal therapies and drug targets: the old, the new and the promising. Cell. Microbiol. 18, 792–799

25. Williamson, P. R. (1994) Biochemical and molecular characterization of the diphenol oxidase of Cryptococcus neoformans: identification as a laccase. J. Bacteriol. 176, 656–664

26. Nosanchuk, J. D., Rudolph, J., Rosas, A. L., and Casadevall, A. (1999) Evidence That Cryptococcus neoformans Is Melanized in Pigeon Excreta: Implications for Pathogenesis. Infect. Immun. 67, 5477 LP – 5479

27. Nosanchuk, J. D., Valadon, P., Feldmesser, M., and Casadevall, A. (1999) Melanization of Cryptococcus neoformans in murine infection. Mol. Cell. Biol. 19, 745–750

28. Nosanchuk, J. D., Rosas, A. L., Lee, S. C., and Casadevall, A. (2000) Melanisation of Cryptococcus neoformans in human brain tissue. Lancet. 355, 2049–2050

29. Chatterjee, S., Prados-Rosales, R., Frases, S., Itin, B., Casadevall, A., and Stark, R. E. (2012) Using solid-state NMR to monitor the molecular consequences of Cryptococcus neoformans melanization with different catecholamine precursors. Biochemistry. 51, 6080–6088

30. Chatterjee, S., Prados-Rosales, R., Itin, B., Casadevall, A., and Stark, R. E. (2015) Solid-state NMR Reveals the Carbon-based Molecular Architecture of Cryptococcus neoformans Fungal Eumelanins in the Cell Wall. J. Biol. Chem. 290, 13779–13790

31. Chatterjee, S., Prados-Rosales, R., Tan, S., Phan, V. C., Chrissian, C., Itin, B., Wang, H., Khajo, A., Magliozzo, R. S., Casadevall, A., and Stark, R. E. (2018) The melanization road more traveled by: Precursor substrate effects on melanin synthesis in cell-free and fungal cell systems. J. Biol. Chem. 293, 20157–20168

32. Nosanchuk, J. D., Stark, R. E., and Casadevall, A. (2015) Fungal Melanin: What do We Know About Structure? Front. Microbiol. 6, 1463

33. Rosas, A. L., Nosanchuk, J. D., Feldmesser, M., Cox, G. M., McDade, H. C., and Casadevall, A. (2000) Synthesis of polymerized melanin by Cryptococcus neoformans in infected rodents. Infect. Immun. 68, 2845–2853

34. Herregodts, P., Michotte, Y., and Ebinger, G. (1989) Regional differences in the distribution of norepinephrine and epinephrine in human cerebral cortex: A neurochemical study using HPLC and electrochemical detection. Neurosci. Lett. 98, 321–326

35. Herregodts, P., Ebinger, G., and Michotte, Y. (1991) Distribution of monoamines in human brain: evidence for neurochemical heterogeneity in subcortical as well as in cortical areas. Brain Res. 542, 300–306

36. Janbon, G., Ormerod, K. L., Paulet, D., Byrnes, E. J. 3rd, Yadav, V., Chatterjee, G., Mullapudi, N., Hon, C.-C., Billmyre, R. B., Brunel, F., Bahn, Y.-S., Chen, W., Chen, Y., Chow, E. W. L., Coppée, J.-Y., Floyd-Averette, A., Gaillardin, C., Gerik, K. J., Goldberg, J., Gonzalez-Hilarion, S., Gujja, S., Hamlin, J. L., Hsueh, Y.-P., Ianiri, G., Jones, S., Kodira, C. D., Kozubowski, L., Lam, W., Marra, M., Mesner, L. D., Mieczkowski, P. A., Moyrand, F., Nielsen, K., Proux, C., Rossignol, T., Schein, J. E., Sun, S., Wollschlaeger, C., Wood, I. A., Zeng, Q., Neuvéglise, C., Newlon, C. S., Perfect, J. R., Lodge, J. K., Idnurm, A., Stajich, J. E., Kronstad, J. W., Sanyal, K., Heitman, J., Fraser, J. A., Cuomo, C. A., and Dietrich, F. S. (2014) Analysis of the genome and transcriptome of Cryptococcus neoformans var. grubii reveals complex RNA expression and microevolution leading to virulence attenuation. PLoS Genet. 10, e1004261

37. Jacobson, E. S., and Emery, H. S. (1991) Temperature regulation of the cryptococcal phenoloxidase. J. Med. Vet. Mycol. bi-monthly Publ. Int. Soc. Hum. Anim. Mycol. 29, 121–124

38. Jacobson, E. S., and Compton, G. M. (1996) Discordant regulation of phenoloxidase and capsular polysaccharide in Cryptococcus neoformans. J. Med. Vet. Mycol. 34, 289–291

39. Garcia-Rivera, J., Eisenman, H. C., Nosanchuk, J. D., Aisen, P., Zaragoza, O., Moadel, T., Dadachova, E., and Casadevall, A. (2005) Comparative analysis of Cryptococcus neoformans acid-resistant particles generated from pigmented cells grown in different laccase substrates. Fungal Genet. Biol. 42, 989–998

40. Chrissian, C., Camacho, E., Fu, M. S., Prados-Rosales, R., Chatterjee, S., Cordero, R. J. B., Lodge, J. K., Casadevall, A., and Stark, R. E. (2020) Melanin deposition in two Cryptococcus species depends on cell-wall composition and flexibility. J. Biol. Chem. 295, 1815–1828

41. Eisenman, H. C., Chow, S.-K., Tsé, K. K., McClelland, E. E., and Casadevall, A. (2011) The effect of L-DOPA on Cryptococcus neoformans growth and gene expression. Virulence. 2, 329–336

42. Wang, Y., and Casadevall, A. (1994) Decreased susceptibility of melanized Cryptococcus neoformans to UV light. Appl. Environ. Microbiol. 60, 3864–3866

43. Polacheck, I., Platt, Y., and Aronovitch, J. (1990) Catecholamines and virulence of Cryptococcus neoformans. Infect. Immun. 58, 2919–2922

44. Meiser, J., Weindl, D., and Hiller, K. (2013) Complexity of dopamine metabolism. Cell Commun. Signal. 11, 34

45. Flierl, M. A., Rittirsch, D., Nadeau, B. A., Chen, A. J., Sarma, J. V., Zetoune, F. S., McGuire, S. R., List, R. P., Day, D. E., Hoesel, L. M., Gao, H., Van Rooijen, N., Huber-Lang, M. S., Neubig, R. R., and Ward, P. A. (2007) Phagocyte-derived catecholamines enhance acute inflammatory injury. Nature. 449, 721–725

46. Lee, S. C., Casadevall, A., and Dickson, D. W. (1996) Immunohistochemical localization of capsular polysaccharide antigen in the central nervous system cells in cryptococcal meningoencephalitis. Am. J. Pathol. 148, 1267–1274

47. Lee, S. C., and Casadevall, A. (1996) Polysaccharide antigen in brain tissue of AIDS patients with cryptococcal meningitis. Clin. Infect. Dis. 23, 194–195

48. Soares, A. R., Marchiosi, R., Siqueira-Soares, R. de C., Barbosa de Lima, R., Dantas dos Santos, W., and Ferrarese-Filho, O. (2014) The role of L-DOPA in plants. Plant Signal. Behav. 9, e28275

49. Folch, J., Lees, M., and Sloane Stanley, G. H. (1957) A simple method for the isolation and purification of total lipides from animal tissues. J. Biol. Chem. 226, 497–509

50. Fung, B. M., Khitrin, A. K., and Ermolaev, K. (2000) An improved broadband decoupling sequence for liquid crystals and solids. J. Magn. Reson. 142, 97–101

51. Melin, V., Henríquez, A., Freer, J., and Contreras, D. (2015) Reactivity of catecholamine-driven Fenton reaction and its relationships with iron(III) speciation. Redox Rep. 20, 89–96

