## Supplemental Figure 1 for "*Cryptococcus neoformans* melanization incorporates multiple catecholamines to produce polytypic melanin"

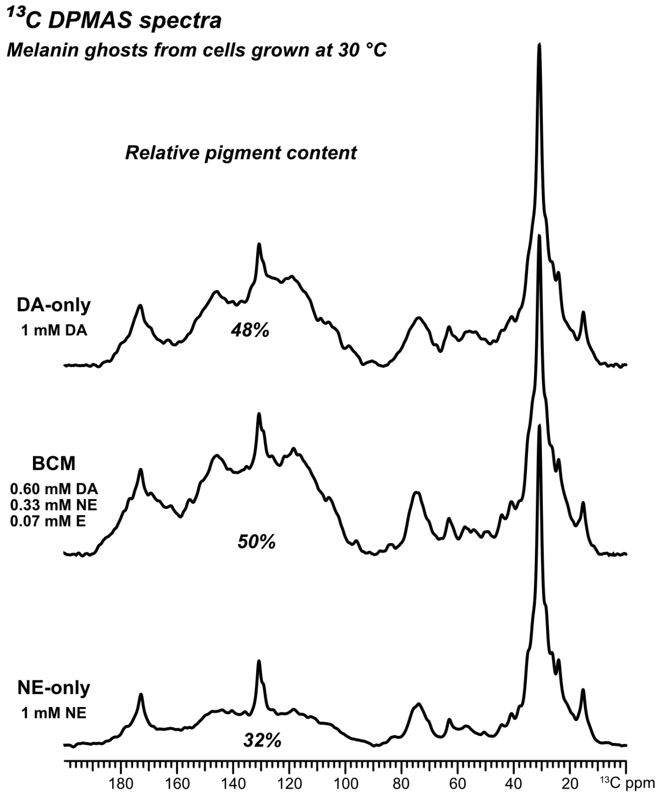


**Figure S1. Quantitatively-reliable solid-state NMR spectra of melanin 'ghosts' from *C. neoformans* cells grown with DA, NE, or BCM.** 1D $^{13}C$ direct-polarization (DPMAS) ssNMR spectra of melanin ghosts isolated from KN99$\alpha$ *C. neoformans* cultures grown for 10 days at 30˚C in minimal media containing either 1 mM dopamine (DA, top), 1 mM norepinephrine (NE, bottom) or in a 'brain catecholamine mixture' of 0.6 mM dopamine, 0.33 mM norepinephrine, and 0.07 mM epinephrine (BCM, middle). The experiments were performed with long (50-s) recycle delays to obtain quantitatively reliable signal intensities that allowed the integration of defined spectral regions using the GNU image manipulation program (GIMP). The relative pigment content was estimated by dividing the integrated area corresponding to the aromatic carbons of the melanin pigment (110-160 ppm) by the total $^{13}C$ NMR signal across the spectrum.
