## Supplemental Figure 2 for "*Cryptococcus neoformans* melanization incorporates multiple catecholamines to produce polytypic melanin"

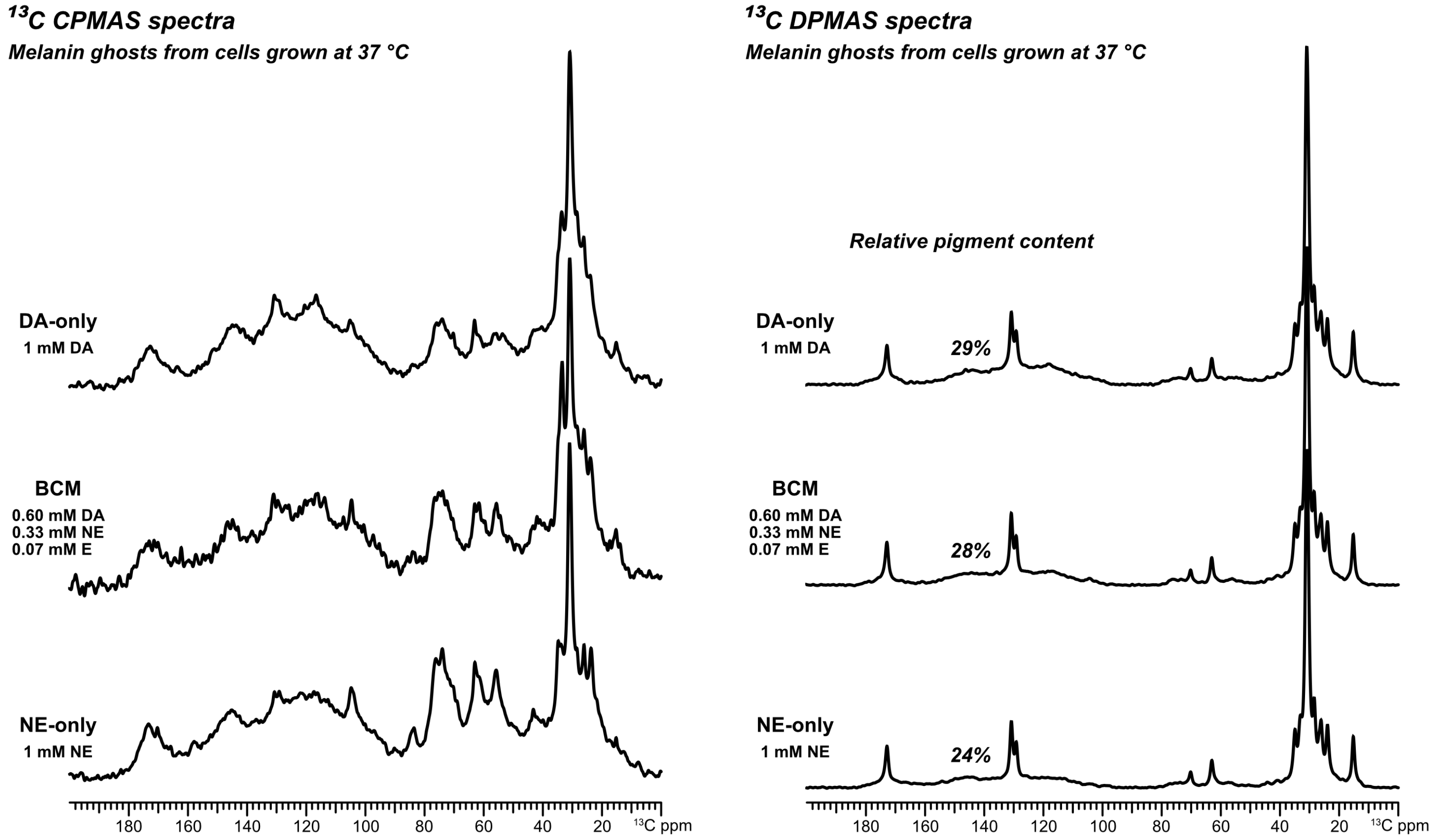


**Figure S2. Solid-state NMR spectra of melanin 'ghosts' from *C. neoformans* cells grown with DA, NE, or BCM.** 1D $^{13}$C cross-polarization (CPMAS) spectra (left) and quantitatively-reliable 1D $^{13}$C direct-polarization spectra (right) of melanin ghosts isolated from KN99$\alpha$ *C. neoformans* cultures grown for 16 days at 37˚C in minimal media containing either 1 mM dopamine (DA, top), 1 mM norepinephrine (NE, bottom) or in a 'brain catecholamine mixture' of 0.6 mM dopamine, 0.33 mM norepinephrine, and 0.07 mM epinephrine (BCM, middle). The DPMAS experiments were performed and the relative pigment content estimated as described in the legend of Figure S1.
